# CD155/CD96 promotes immunosuppression in lung adenocarcinoma (LUAD)

**DOI:** 10.1101/688812

**Authors:** Weiling He, Hui Zhang, Shuhua Li, Yongmei Cui, Ying Zhu, Junfeng Zhu, Yiyan Lei, Run Lin, Di Xu, Zheng Zhu, Wenting Jiang, Han Wang, Zunfu Ke

## Abstract

Lung adenocarcinoma (LUAD) remains one of the leading causes of death in patients with cancer. The association of CD155 with CD96 transmits an inhibitory signal and suppresses antitumor immune response. This study investigates the effect of CD155/CD96 on immune suppression in LUAD. We demonstrate that LUAD patients with high CD155 expression suffer from immune suppression and experience a poor prognosis, which coincides with an inhibited AKT-mTOR signaling pathway in CD8 T cells and subsequently up-regulated CD96 expression. Moreover, the inhibition effect can be reversed by CD96 blocking antibody. High CD155 expression inhibited the release of IFNγ from CD8 cells. Moreover, Blocking CD96 restored IFNγ production in CD8 T cells and neutralized the inhibition of IFNγ production in CD8 T cells mediated by CD155. Animal experiments showed that CD155-mediated LUAD growth might depend on its suppression antitumor immune response in the tumor microenvironment in PDX mice. In conclusion, our results suggest that LUAD cells suppress antitumor immune response in the tumor microenvironment through CD155/CD96. CD155/CD96 could be a potential therapeutic target for LUAD patients.

**Abbreviations:** LUAD: lung adenocarcinoma; IFNγ: interferon gamma; PDX: patient-derived xenograft; NSCLC: non-small cell lung cancer; PRR: poliovirus receptor–related; MDSCs: myeloid-derived suppressor cells; PRR: poliovirus receptor–related; STR: short tandem repeat; IRS: immunoreactive score; SI: staining intensity; PP: percentage of positive cells; RT-PCR: reverse transcription-polymerase chain reaction; PBS: phosphate-buffered saline; PBMCs: peripheral blood mononuclear cells; SDS–PAGE: sodium dodecyl sulfate-polyacrylamide gel electrophoresis; rCD155: recombinant human CD155; LUAD cells: lung adenocarcinoma cells; TILs: tumor-infiltrating lymphocytes; GzmB: granzyme B; IL-2 (Interleukin-2); TNF-α : tumor necrosis factor-alpha; PI: propidium Iodide; PDX: patient-derived xenograft; TIGIT: T cell immunoreceptor with Igand ITIM domains; WBC: white blood cells; MFI: mean fluorescence intensity; HPF: high power field

## Introduction

Lung cancer is the leading cause of death among cancer patients. More than 1 million deaths are related to lung cancer annually (Ding *et al*, 2008), and approximately 1.2 million new lung cancer cases are diagnosed each year (Jemal *et al*, 2011). Lung adenocarcinoma (LUAD) is the most common type of non-small cell lung cancer (NSCLC). Surgery remains the first-line treatment and the most successful option in located disease of LUAD (Molina *et al*, 2008). Despite a decline in the cancer death rate over the past two decades, LUAD has a 5-year survival rate of 10-15% in stage IV due to its late-stage diagnosis and lack of effective therapeutic options (Imielinski *et al*, 2012).

Immune escape represents the failure of the immune system in preventing carcinogenesis (Kim *et al*, 2007; Swann & Smyth, 2007). The tumor microenvironment is composed of regulatory T cells, tumor-associated macrophages and myeloid-derived suppressor cells (MDSCs) (Noy & Pollard, 2014; Sato *et al*, 2005; Talmadge & Gabrilovich, 2013), which are the signature of chronic inflammation and immune suppression in the tumor milieu (Crespo *et al*, 2013). Recently, the development of cancer immunotherapy that uses tumor-targeted monoclonal antibodies has achieved broad therapeutic efficacy (Brahmer *et al*, 2012). The application of monoclonal antibody against PD-1/PD-L1 was associated with longer progression-free and overall survival and fewer treatment-related adverse events than was platinum-based combination chemotherapy in patients with previously untreated advanced NSCLC (Herbst *et al*, 2016; Reck *et al*, 2016). Given the only 20-30% response rate of lung cancer when targeting PD-1/PD-L1, we speculated that other molecules or mechanisms might be involved in limiting the application of current immunotherapies for lung cancer patients.

CD155, a member of poliovirus receptor–related (PRR) family, was initially identified as a receptor for poliovirus in humans (Mendelsohn *et al*, 1989). The CD155 transcript is ubiquitously expressed in humans and mice (Koike *et al*, 1990; Mendelsohn *et al*, 1989). Recently, CD155 has been identified as the ligand of the Ig-like receptors Tactile (CD96) and DNAM-1 (CD226) on T cells. The interaction of CD155 with CD96 transmits an inhibitory signal. The interaction of CD155 with CD226 enhances the immune response (Bottino *et al*, 2003; Shibuya *et al*, 1996; Yu *et al*, 2009). Expression of CD155 has been shown to be elevated in many types of tumors (Brooks *et al*, 2017; Iguchi-Manaka *et al*, 2016) and is involved with immune suppression in melanoma (Inozume *et al*, 2016). Recently, CD96^−/−^ mice displayed hyper-inflammatory responses, resulting in resistance to carcinogenesis and experimental lung metastases (Chan *et al*, 2014). Blocking CD96 can improve tumor control in mice (Blake *et al*, 2016). However, how CD155/CD96 is involved in immune response in the tumor microenvironment of LUAD remains unknown.

In the present study, we found that CD155 expression was increased in tumor tissue and was associated with immune suppression in the tumor microenvironment of LUAD. Blocking CD155/CD96 interaction restores CD8 T cell effector functions by reversing CD8 T cell exhaustion, suggesting a possible therapeutic role of CD155/CD96 in fighting LUAD.

## Results

### CD155 expression was associated with immune suppression in the tumor microenvironment of LUAD

To investigate CD155 expression characteristics in LUAD tissues, we first performed western blotting to compare its expression between tumor and para-tumor lung tissues. We found that CD155 expression was substantially stronger in LUAD tissues than that in para-tumor normal lung tissues (Figure 1A). The increased CD155 expression in LUAD tissues was confirmed by IHC (Figure 1B). Lung adenocarcinoma cells (LUADCs) were isolated from tumor tissues in 6 CD155^high^ and 6 CD155^low^ LUAD patients using NanoVelcro as described previously (Lin *et al*, 2014) (Figure S1). The primary LUADCs isolated from tumor tissues were also CD155 positive as measured by immunofluorescence (Figure 1C). Further, CD155 high expression in tumor tissues predicted a poor prognosis in LUAD (Figure S2). CD155, as the molecule of a co-inhibitory pathway, suppresses the anti-melanoma immune response (Inozume *et al*, 2016). We speculate that the low survival rate in CD155^high^ patients might be due to the immune suppression in the tumor microenvironment mediated by CD155.

**Figure 1.**
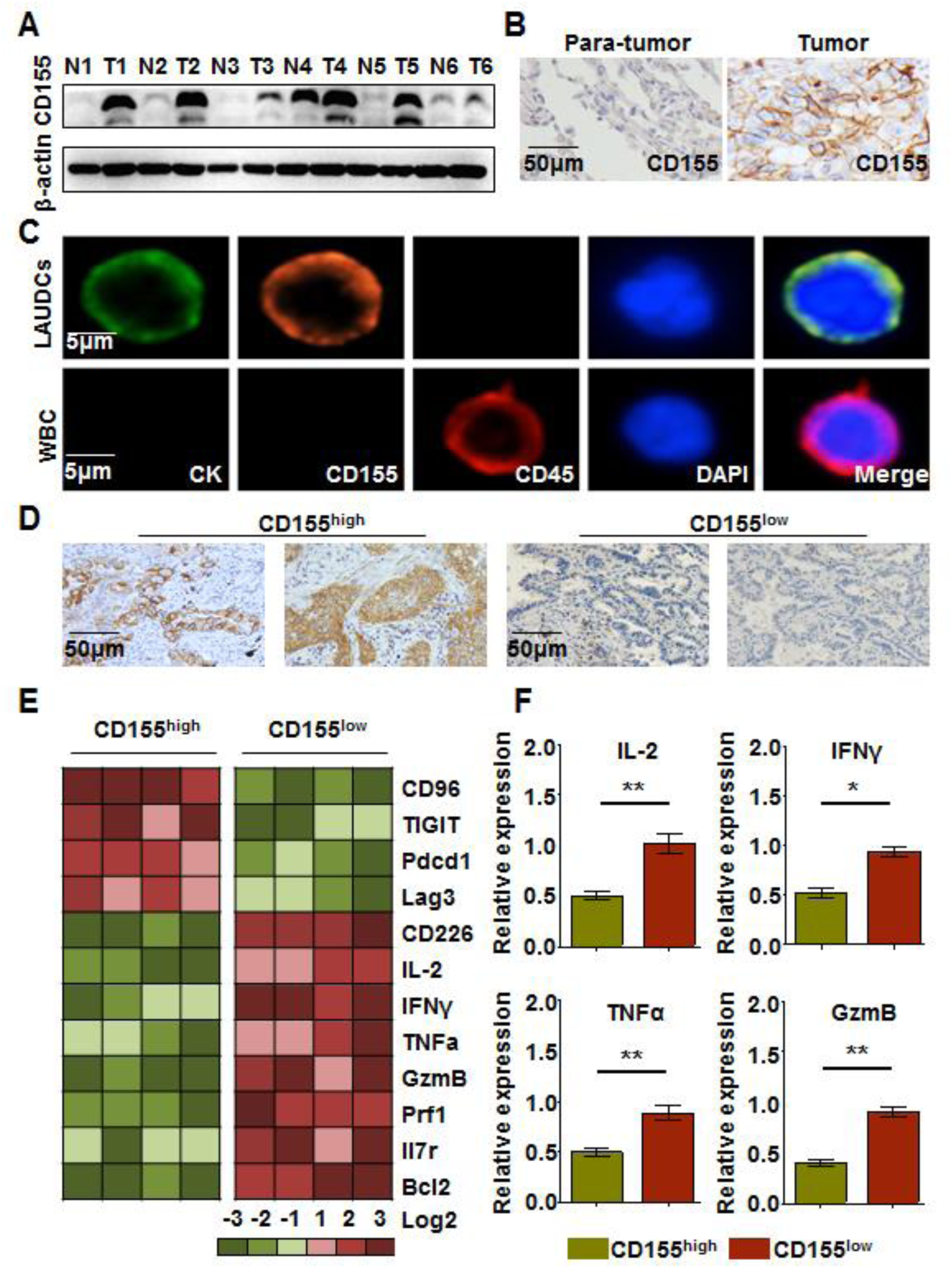
CD155 expression and immune suppression in the tumor microenvironment of LUAD. (A), CD155 expression in tumor or para-tumor lung tissues from patients (n=6) with LUAD was measured by western blotting. All of 6 representative patients were considered CD155^high^. (B), CD155 expression in tumor or para-tumor lung tissues was detected by immunohistochemistry (200×). (C), Primary LUADCs were isolated from 6 CD155^high^ patients and expanded as in Supplementary Figure 1. CD155 expression in primary LUADCs was detected by immunofluorescence. White blood cells (WBC) were used as control. (D), CD155^high^ or CD155^low^ expression was determined by immunohistochemistry (200×). (E), RNA was extracted from tumor tissue. Gene expression was measured by RT-PCR and shown as a heat map. (F), Gene expression of IL-2, IFNγ, TNFα and granzyme B (GzmB) was summarized from 24 independent samples (13 for CD155^high^ and 11 for CD155^low^). Error bars show SEM. The data are shown as the mean ± SEM, *p<0.05, **p<0.01.

To further explore the involvement of CD155 in the antitumor immune response in LUAD, we first studied the association of CD155 with the immune response in the tumor microenvironment of LUAD. CD155^high^ and CD155^low^ were identified according to its expression intensity as described in the methods section (Figure 1D). We found that the transcripts of inhibitory molecules, such as CD96, TIGIT, Pdcd1 and Lag-3, were substantially higher in CD155^high^ patients, while CD226 was lower in CD155^high^ patients (Figure 1E and S3). In contrast, gene expression of T cell effector function-associated molecules was essentially lower in 13 CD155^high^ patients compared with that in 11 CD155^low^ patients (Figure 1E-F). Thus, high expression of CD155 in LUAD might be associated with immune suppression in tumor microenvironment.

### LUADCs impaired the effector functions of CD8 T cells through direct cell-cell contact

Tumor-infiltrating lymphocytes (TILs) predict a better prognosis in patients with colorectal cancer (Pages *et al*, 2005). Here, we showed that a high density of CD8 T cell infiltration was associated with better survival in patients with LUAD (Figure S4), which was in accordance with a previous report (Kawai *et al*, 2008). CD8 T cells are the effector cells in antitumor immune response. However, TILs are functionally exhausted in the tumor microenvironment, which has not fully been understood yet. To elucidate how cancer cells influence CD8 T cells in the tumor microenvironment, we first performed a cell-cell contact co-culture of CD8 T cells (from the same CD155^high^ LUAD patient) and LUADCs (from 6 independent CD155^high^ patients) in vitro (Figure 2A). We found that GzmB (granzyme B) and Perforin expression on CD8 T cells was inhibited, as measured by flow cytometry, when co-cultured with LUADCs (Figure 2B-E). Further, production of IL-2 (Interleukin-2), TNFα (tumor necrosis factor-alpha) and IFNγ cytokines in CD8 T cells was also suppressed (Figure 2F-I). However, cytokine production was not affected when we separately cultured the T cells (from the same CD155^high^ LUAD patient) and LUADCs (from 6 independent CD155^high^ patients) using a cell culture insert with a 0.4-μm pore size (Figure 2J-K). Thus, cancer cells impaired CD8 T cell effector function through cell-cell contact, which might contribute to immune suppression in the tumor microenvironment of LUAD.

**Figure 2.**
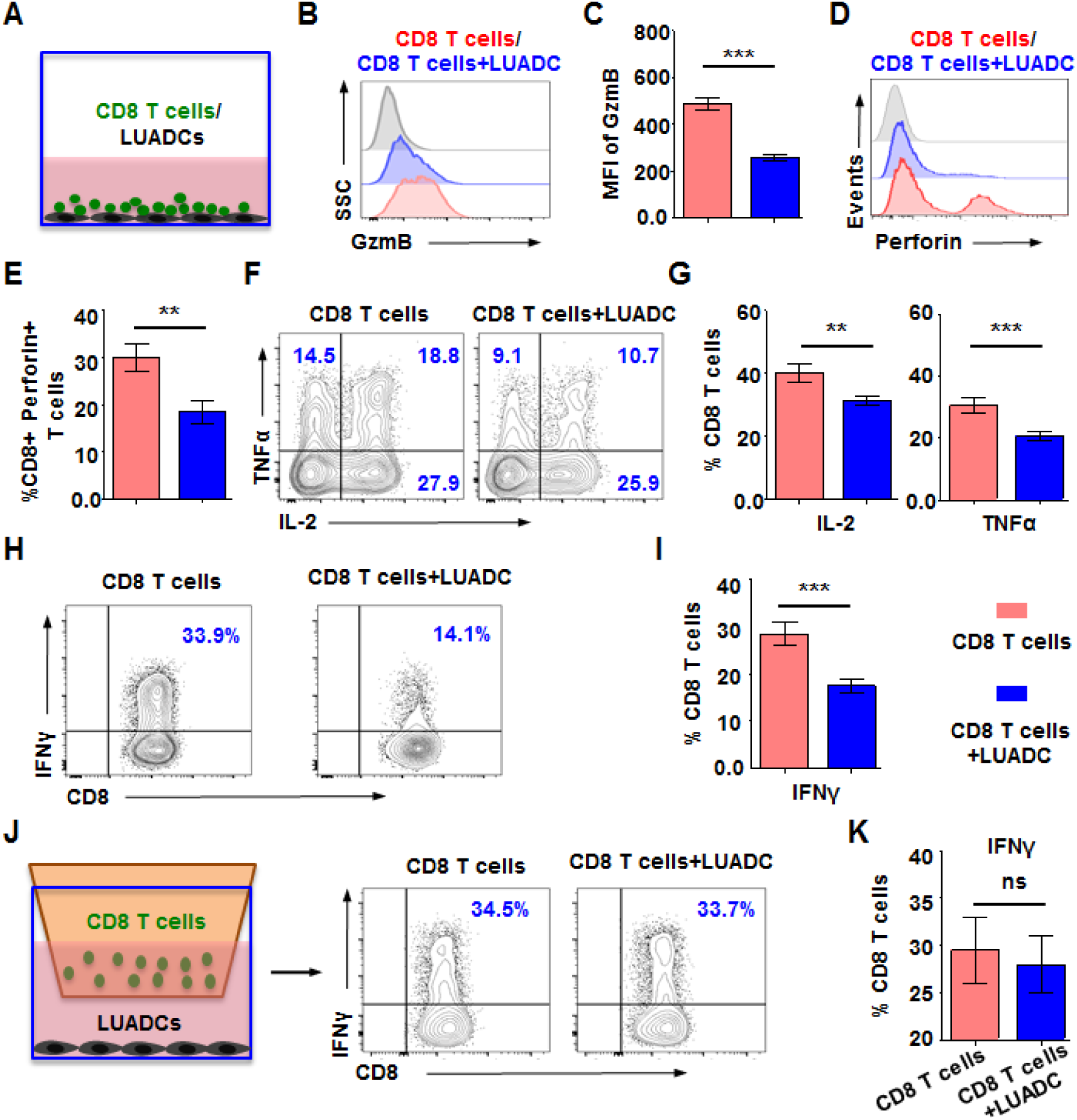
LUAD cells impaired CD8 T cell effector functions. (A), Scheme of CD8 T cells (from the matched CD155^high^ LUAD patient) co-cultured with LUADC1-6 from 6 independent CD155^high^ patients in a cell-cell contact manner. CD8 T cells were stimulated with beads for 3 days. Representative results are shown as follows (B-I). (B), GzmB expression on CD8 T cells was analyzed by flow cytometry. (C), Mean fluorescence intensity (MFI) of GzmB in CD8 T cells was summarized. (D), Perforin expression on CD8 T cells was analyzed by flow cytometry. (E), Percentage of Perforin-expressing CD8 T cells was summarized. (F), IL-2 and TNFα production in CD8 T cells was measured by flow cytometry. (G), Percentages of IL-2- and TNFα-producing CD8 T cells were summarized. (H), IFNγ production in CD8 T cells was measured by flow cytometry. (I), Percentage of IFNγ+ CD8 T cells was summarized. (J), Scheme of CD8 T cells (from the same CD155^high^ LUAD patient) co-cultured separately with LUADC1-6 from 6 independent CD155^high^ patients. A cell culture insert with 0.4-μm pore size was used to culture CD8 T cells and tumor cells separately. IFNγ production in CD8 T cells was measured by flow cytometry. Representative results are shown as follows (K). (K), Percentage of IFNγ+ CD8 T cells was summarized. The data are shown as the mean±SEM of 3 independent experiments, **p<0.01 ***p<0.001. ns: not significant.

### LUADCs suppressed CD8 T cell function through CD155

CD155 expression in the melanoma correlated with immune suppression in the tumor microenvironment ^[21]^. To confirm the hypothesis that CD155 might be involved in T cell inhibition mediated by LUADCs, we first determined CD155 expression in LUADC1-6 by western blotting. Compared with CD155-low level in BEAS-2B cells, LUADC1-6 cells appear CD155-high (Figure 3A). CD155 expression in LUADCs was further confirmed by flow cytometry (Figure 3B). In a T cell-cancer cell co-culture system (Figure 3C), LUADCs decreased the expression of p-AKT and p-mTOR in CD8 T cells, as measured by flow cytometry. The phosphorylation of S6K and 4EBP1 in CD8 T cells was also decreased by cancer cells (Figure 3D). Downregulation of CD155 in LUADCs by RNAi abolished the inhibition on CD8 T cells. The suppression of AKT, mTOR, S6K and 4EBP1 in CD8 T cells was reversed by knocking down CD155 in LUADCs (Figure 3D). These data demonstrated that LUADCs suppress CD8 T cell effector function, which could lead to poor antitumor immune response. Further, IFNγ production in CD8 T cells was decreased when co-cultured with LUADCs. Knocking down CD155 in cancer cells could reverse the inhibition (Figure 3E-F). Additionally, CD155 upregulation in LUADCs further suppressed IFNγ production in CD8 T cells compared with that in LUADC-vector cells (Figure 3G-H). In summary, CD155 mediated LUADCs’ suppression on CD8 T cell function.

**Figure 3.**
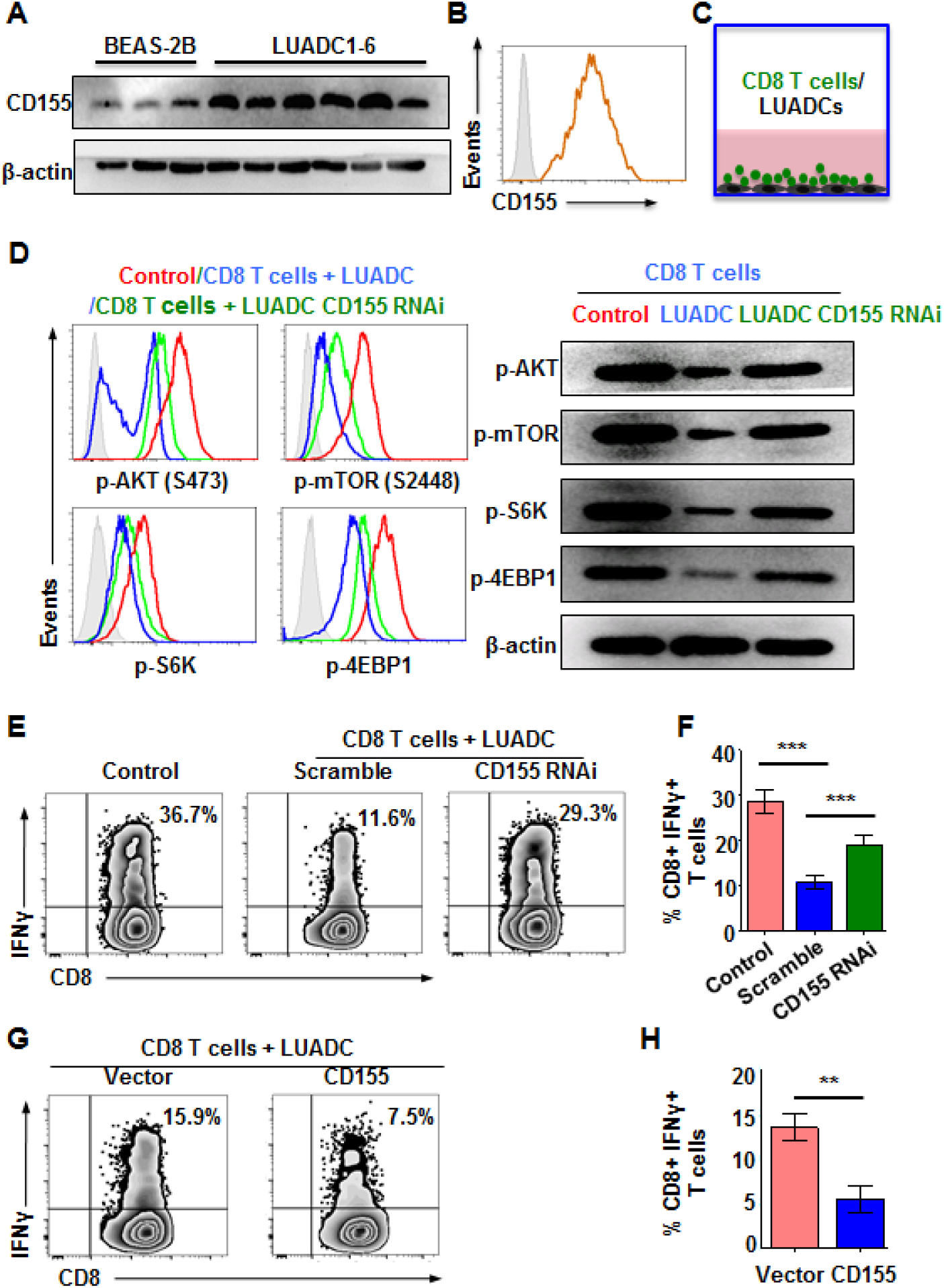
LUAD cells suppress CD8+ T cell effector functions through CD155. Representative results are shown as follows (A-H). (A), CD155 expression in LUADC1-6 cells (from 6 independent CD155^high^ patients) was detected by western blotting. (B), CD155 expression in LUADC was detected by flow cytometry. (C), CD8 T cells (from the matched CD155^high^ LUAD patient) were stimulated with anti-CD3/CD28 beads and co-cultured with LUADC. (D) CD8 T cells were co-cultured with LUAD cells that were treated with Scramble or CD155 RNAi. CD8^+^ T cells cultured alone was used as control. Phosphorylation of AKT, mTOR, S6K and 4EBP1 in CD8^+^ T cells was measured by flow cytometry and immunoblot, respectively. (E), CD8 T cells (from the matched CD155^high^ LUAD patient) were co-cultured with or without LUADC cells that were treated with CD155-specific or scramble RNAi for 3 d. CD8^+^ T cells cultured alone was used as control. IFNγ production in CD8 T cells was measured by flow cytometry and summarized in F. (G), CD8 T cells (from the matched CD155^high^ LUAD patient) were co-cultured with LUADC in which CD155 was stably upregulated for 3 d. IFNγ production in CD8 T cells was measured by flow cytometry and summarized in H. The data are shown as the mean±SEM of 3 independent experiments, **p<0.01, ***p<0.001.

### CD155-independent tumor growth in LUADCs

CD155 was stably downregulated in LUADCs (from 6 independent CD155^high^ patients) or overexpressed in LUADCs (from 6 independent CD155^low^ patients). Knockdown or overexpression was confirmed by western blotting and flow cytometry (Figure S5). We found that neither knockdown nor overexpression of CD155 affected cancer cell proliferation, as measured by cell number (Figure S6A-B). We used PI (Propidium Iodide) to investigate the cell cycle by flow cytometry and observed that CD155 knockdown did not change the cell cycle distribution of cancer cells. Further, CD155 overexpression showed no effects on the cell cycle (Figure S6C-E). To study whether tumor growth was affected by CD155 in vivo, we utilized non-invasive imaging. LUADCs were stably transfected with luciferase and inoculated into NOG mice subcutaneously. The data showed no difference in tumor growth between mice that received LUADCs-vector or LUADCs-CD155 cells (Figure S6F-G).

### CD155 promotes tumor growth in LUAD tumor-bearing mice by impairing the antitumor immune response

The tumor microenvironment is infiltrated with tumor-associated macrophages, myeloid-derived suppressor cells and regulatory T cells (Hanahan & Weinberg, 2011). The stromal cells also play critical roles in shaping the tumor microenvironment (Hanahan & Coussens, 2012). To better duplicate the tumor microenvironment similar to LUAD patients, we used a PDX mouse model as described in the method section to study the immune response in the tumor microenvironment. Previous study shows that CD155/96 is important for NK cells in antitumor immune response. In this mouse model, the CD56+ NK cells were sorted out from the injected PBMC (from 6 independent CD155^low^ patients), excluding the antitumor effects by NK cells. To investigate whether CD155/CD96 affects immune reaction and tumor growth, PDX mice were treated with rCD155 or vehicle (Figure 4A). We first confirmed the binding of rCD155 to CD96 and that decreased IFNγ production in CD8 T cells (Figure S7). The infiltration of CD8 T cells in the tumor microenvironment was decreased to some extent by rCD155 treatment as measured by IHC (Figure 4B-C). To measure the effector function of the tumor infiltrated CD8 T cells, CD8 T cells were isolated from the tumor tissue and measured IFNγ production by flow cytometry. rCD155 treatment decreased IFNγ production in CD8 T cells isolated from tumor tissue (Figure 4D-E). The IFNγ transcript was also decreased by rCD155 treatment as measured by RT-PCR (Figure 4F). To further understand the functional status of CD8 T cells in the tumor microenvironment, we measured TNF-α, GzmB and Perforin expression in the tumor microenvironment by RT-PCR. We found that rCD155 treatment decreased TNF-α, GzmB and Perforin expression in the tumor microenvironment (Figure 4G). Importantly, rCD155 treatment promoted tumor growth in the PDX mice (Figure 4H).

**Figure 4.**
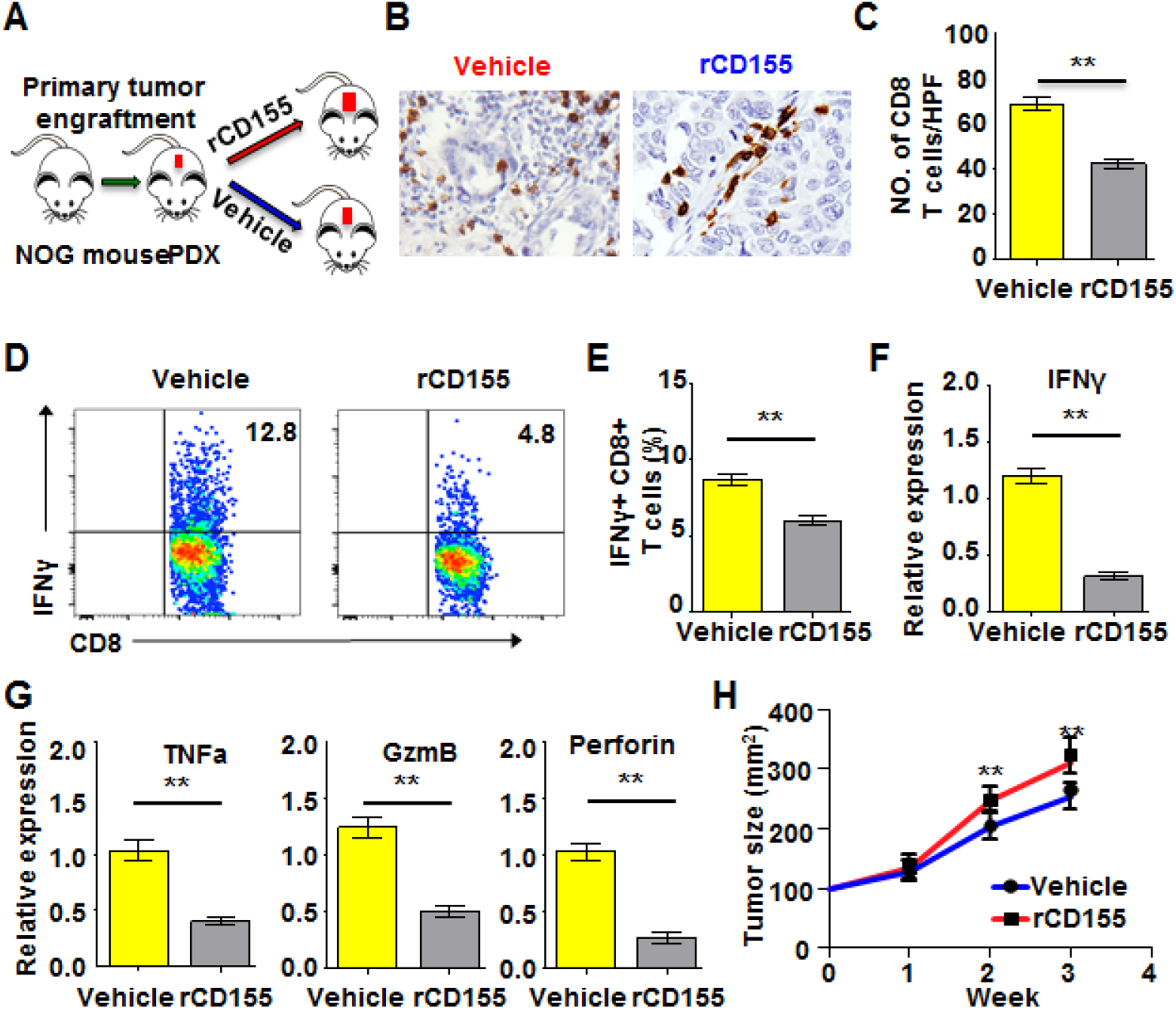
CD155 suppressed the immune response in the tumor microenvironment and promoted tumor growth. (A), Scheme of the patient-derived xenograft (PDX) experiment. Fresh tumor samples (from 6 independent CD155^low^ patients) were collected and engrafted into NOG mice subcutaneously. One week later, CD56^−^ PBMC from the matched donors were injected intraperitoneally for immune reconstitution. PDX mice were treated with recombinant CD155 (rCD155, 5mg/kg) or vehicle. (B), The number of CD8 T cells in the tumor microenvironment was measured by IHC (400×). (C) The number of CD8 T cells per high power field (HPF) was summarized from 6 independent samples. (D), Tumor tissue was digested to generate a single-cell suspension. IFNγ production in CD8 T cells was measured by flow cytometry. Representative flow plots were gated on CD8 T cells. (E), Percentages of CD8^+^IFNγ^+^ cells were summarized from 6 independent samples. (F, G), RNA was extracted from tumor tissue. Transcripts of IFNγ, TNFα, GzmB and Perforin expression in the tissue was measured by RT-PCR. The data were summarized from 6 independent samples. (H), Mice were treated with rCD155 or vector, and tumor growth was monitored. The data are shown as the mean±SEM, **p<0.01. Ns: not significant.

To further investigate the function of CD155 on immune response in the tumor microenvironment of LUAD, CD155 was stably overexpressed in LUADC cells (from 6 independent CD155^low^ patients). Cells were inoculated into NOG mice subcutaneously. The mice were subsequently reconstituted with or without human PBMCs. We found that tumors in NOG mice that lacked human PBMC reconstitution grew substantially faster than those in mice carrying a human immune system (Figure S8A-B). Further, in mice reconstituted with human PBMCs, tumor growth was substantially faster in mice that received LUADC-CD155 cells than in those that received LUADC-vector cells (Figure S8C-D). Moreover, mice that received LUADC-CD155 cells showed a poorer prognosis than those that received LUADC-vector cells (Figure S8E).

Therefore, these results demonstrated that CD155-mediated LUAD growth might depend on its suppression antitumor immune response in the tumor microenvironment.

### Dysregulation of CD96/CD226 identified reduced CD8 T cell effector functions in LUAD

CD96, as a co-inhibitory molecule, competes with the co-stimulatory molecule of CD226 for binding to CD155. The lost balance between these two molecules might lead to a hypo- or hyper-immune response. We found that CD8 T cells from LUAD patients expressed higher levels of CD96 compared with healthy controls (HC). CD96 expression was even higher in TILs than in T cells from the circulation (Figure 5A-B). CD226 expression was substantially lower in T cells from LUAD patients than that in HC. CD226 expression in TILs was further decreased compared with that from the circulation (Figure 5C-D). The phosphorylation of AKT and mTOR was lower in CD96^+^ CD8 T cells from LUAD patients than that in CD96^−^ CD8 T cells. The phosphorylation of mTOR downstream molecules also decreased in CD96^+^ cells compared with that in CD96^−^ cells, as measured by flow cytometry and western blotting (Figure 5E). To compare the effector functions of CD96+ and CD96-CD8 T cells, intracellular cytokine production was measured by flow cytometry. IFNγ and TNFα production in CD96+ and CD96^−^ T cells was confirmed by flow cytometry (Figure 5F-G). CD96 expression in CD8 T cells from LUAD patients was associated with decreased T cell effector functions. The increased CD96 expression in CD8 T cells from LUAD patients might be closely associated with the poor immune response in tumor microenvironment.

**Figure 5.**
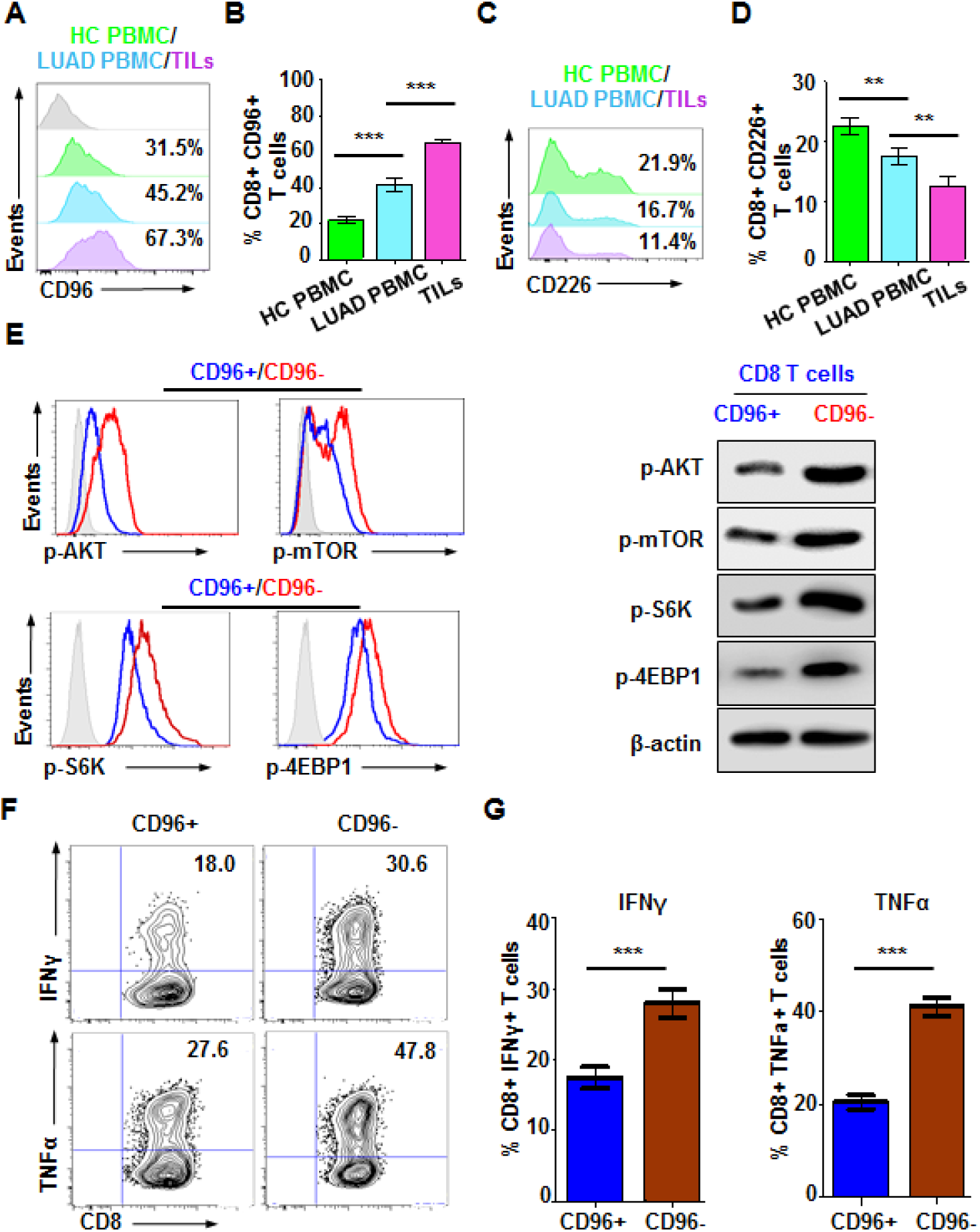
Dysregulation of CD96/CD226 identified CD8 T cell exhaustion in LUAD. (A-D), CD96 or CD226 expression on CD8 T cells from PBMCs of 24 matched patients with LUAD (13 CD155^high^ and 11 CD155^low^) or healthy controls (HC) (n=24) and from tumor-infiltrating lymphocytes (TILs) was analyzed by flow cytometry. Percentages of CD8^+^CD96^+^ or CD8^+^CD226^+^ cells in PBMCs (n=24) or TILs (n=24) were summarized and shown as bar graphs. Error bars show SEM. (E), CD8^+^CD96^+^ or CD8^+^96^−^ cells were sorted from PBMCs in LUAD patient. Cells were stimulated with anti-CD3/CD28 beads. Phosphorylation of AKT, mTOR S6K and 4EBP1 was measured by cytometry and immunoblot, respectively. (F, G), TNFα and IFNγ production in CD96^+^ or CD96^−^ CD8 T cells from LUAD patient was measured by flow cytometry. The data are shown as the mean±SEM, **p<0.01, ***p<0.001.

### LUAD cells suppress effector functions of CD8 T cell through CD155/CD96

The tumor microenvironment is infiltrated with exhausted T cells (Crespo *et al*, 2013; Jiang *et al*, 2015). CD96, a recently identified co-inhibitory molecule that competes with CD226 for the ligand CD155, was increased on CD8 T cells in the LUAD tumor microenvironment (Chan *et al*, 2014). To determine how CD96 expression on CD8 T cells is regulated in the tumor microenvironment by cancer cells, we performed cell-cell contact T cell-cancer cell co-culture as described above and found that CD96 expression on CD8 T cells was increased when co-cultured with CD155^high^ cancer cells (from 6 independent CD155^high^ patients) (Figure 6A), whereas the co-stimulatory receptor CD226 was inhibited by CD155^high^ cancer cells (Figure 6B). As shown in Figure 3D, AKT-mTOR pathway in CD8 T cells was inhibited by culturing with CD155^high^ cancer cells. We found that blocking CD96 can rescue the inhibition. The phosphorylation of AKT-mTOR was increased by blocking CD96 as measured by western blot (Figure 6C). Also, the phosphorylation of 6SK and 4EBP1, which are downstream molecules of mTOR, was also increased in CD8 T cells when CD96 was blocked, as measured by flow cytometry (Figure 6D). Blocking CD96 in the T cell-cancer cell co-culture system increased IFNγ production in CD8 T cells (Figure 6E). IFNγ production in CD8 T cells was decreased when co-cultured with CD155^high^ LUAD cells, which was further decreased when CD155 was overexpressed in CD155^high^ LUAD cells (Figure 6F). However, CD96 blockade could neutralize the inhibition mediated by CD155 expression (Figure 6F). Thus, LUAD cells suppress the T cell response through CD155/CD96 signaling in the tumor microenvironment.

**Figure 6.**
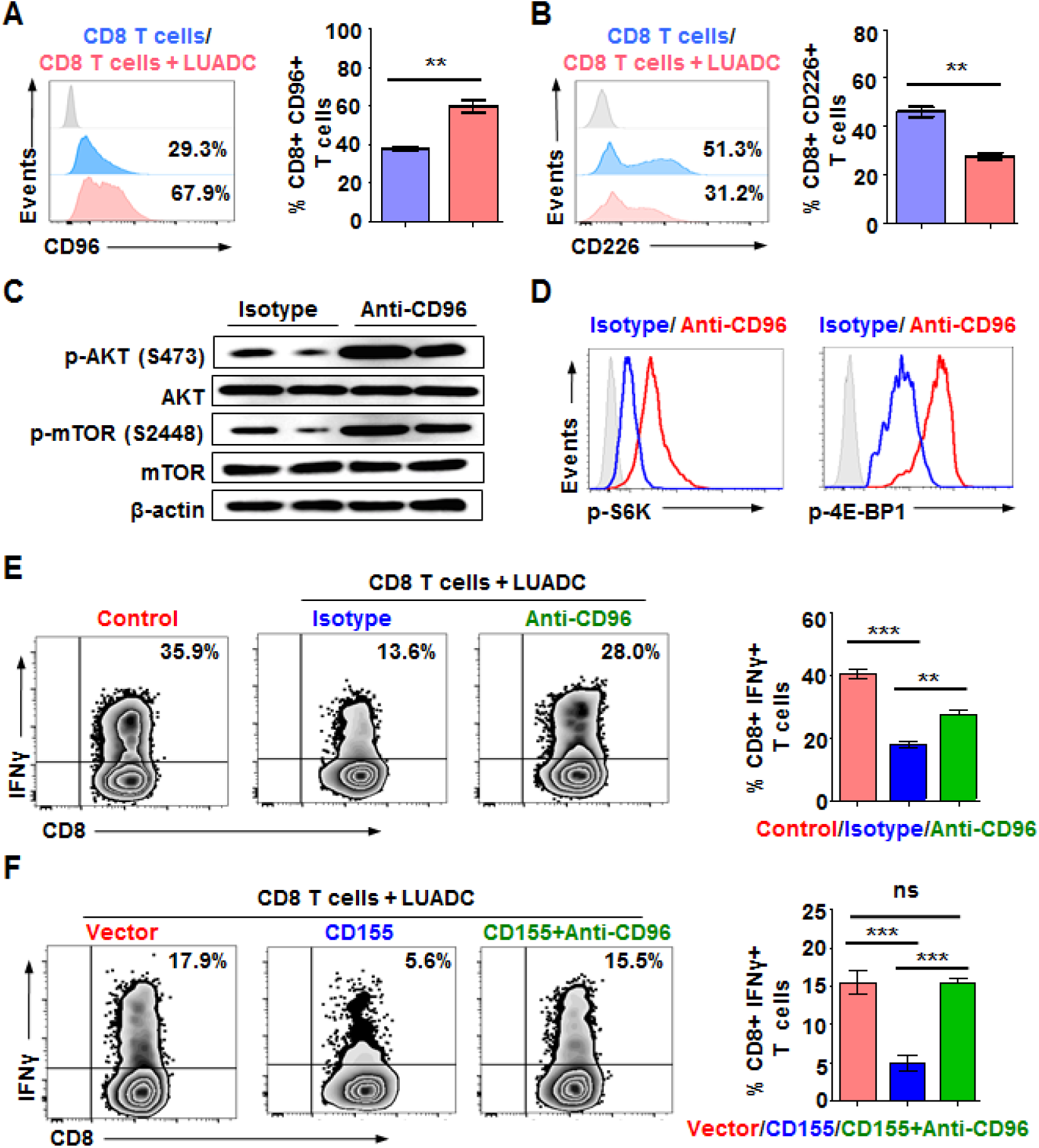
LUAD cells suppressed anti-tumor immune response through the CD155/CD96 pathway. (A, B), CD8 T cells isolated from the matched LUAD patient were stimulated with anti-CD3/CD28 beads and co-cultured with LUADC1 from the same patien for 3 d. CD96 and CD226 expression on CD8 T cells was measured by flow cytometry. Percentage of CD8^+^CD96^+^ or CD8^+^CD226^+^ T cells was summarized from 8 independent samples. (C), Phosphorylation of AKT and mTOR in CD8 T cells were measured by western blotting. The expression of p-S6K and p-4EBP1 was determined by flow cytometry. (D, E), CD8^+^ T cells were co-cultured with autologous LUAD cells. Anti-CD96 antibody (5 μg/ml) was added in some experiments. CD8 T cells cultured alone was used as control. IFNγ production in CD8 T cells was measured by flow cytometry. Percentage of IFNγ^+^ CD8 T cells was summarized from 6 independent samples (right panel). (F) CD155 or empty vector was transfected into LUADC1. CD8 T cells were co-cultured with LUADC1-CD155 or LUADC1-vector for 3 d. Anti-CD96 antibody (5 μg/ml) was added in the experiments. IFNγ production in CD8 T cells was measured by flow cytometry. The percentage of IFNγ^+^ CD8 T cells was summarized from 6 independent samples (right panel). The data are shown as the mean±SEM, **p<0.01, ***p<0.001. ns: not significant.

## Discussion

In the tumor microenvironment, the upregulation of negative signals on T-cell responses suppresses the antitumor immune response and promotes tumor progression (Zou & Chen, 2008). In this study, we showed that CD155 expression was increased in LUAD tumor tissue, and LUAD cells suppressed the immune response in the tumor microenvironment through CD155/CD96. However, CD155 itself showed no effects on cancer cell proliferation in vitro or tumor growth in vivo.

CD8 T cells are the effector cells in the antitumor immune response in most tumor models (Rosenberg *et al*, 2004). However, T cells that have infiltrated into the tumor microenvironment are exhausted and functionally impaired (Ahmadzadeh *et al*, 2009; Jiang *et al*, 2015). In the current study, we found that LUAD cells impaired the effector functions of CD8 T cells through cell-cell contact, which explained the immune suppression in the tumor microenvironment. In addition to tumor-associated macrophages, MDSC and regulatory T cells, cancer cells can impair CD8 T cell functions directly and escape immune attack.

CD155 functions as a cell adhesion molecule that enhances glioma cell migration (Sloan *et al*, 2005), and its expression is increased at the RNA and protein levels in tumor tissues (Chandramohan *et al*, 2017; Masson *et al*, 2001). The association of CD155 to TIGIT (T cell immunoreceptor with Ig and ITIM domains) transmits a negative signal to T cells and suppresses immune response (He *et al*, 2017). In the current study, we found that CD155 expression was significantly increased in LUAD tissue. The primary LUAD cells isolated from patients with LUAD expressed high levels of CD155, which correlated with immune suppression in the tumor microenvironment. The CD155-associated hypo-immune response in the tumor microenvironment of LUAD could lead to immune invasion and promote tumor growth. In addition, CD155^high^ patients showed poorer survival than CD155^low^ patients, which was in accordance with a previous report (Atsumi *et al*, 2013). Interestingly, we found that CD155 expression did not affect cancer cell proliferation in vitro or tumor growth in vivo. Further, tumor growth was greater in NOG mice without a human immune system than in mice with a reconstituted human immune system. CD155 overexpression in cancer cells resulted in greater tumor growth and poorer survival in tumor-bearing NOG mice reconstituted with human immune system. CD155 promoted tumor growth in mice reconstituted with the human system through limiting the immune response in the tumor microenvironment. These data further suggest that CD155 mediates LUAD progression by manipulating the immune system, which is independent of cancer cell proliferation.

To better understand how CD155 regulates the immune response mediated by CD8 T cell in the tumor microenvironment, we established a PDX model using patient-derived tumor tissues and reconstituted the mice with PBMC from the same donors. To exclude the anti-tumor effect mediated by CD56+ NK cells, CD56+ NK cells in PBMC were sorted out before injection. We found that rCD155 treatment decreased CD8 T cell in the tumor microenvironment and IFNγ production in the infiltrated CD8 T cells was decreased by rCD155. In addition, rCD155 decreased the transcripts of IFNγ, TNF-α, GzmB and Perforin in the tumor tissue. The suppressed immune response contributed to tumor growth in the PDX mice. Thus, enhancing the antitumor immune response targeting CD155 in the tumor microenvironment could be a good therapeutic strategic for LUAD.

The balance between the inhibitory signal CD96/CD155 and the co-stimulatory signal CD226/CD155 is important for immune homeostasis (Chan *et al*, 2014; Gao *et al*, 2017). CD96 blockade inhibits experimental metastases (Blake *et al*, 2016) and prevents tumor from relapsing in a transgenic pancreatic ductal adenocarcinoma mouse model (Brooks *et al*, 2017). In the current study, we found that CD96 expression was increased in CD8 T cells from LUAD patients. CD226, the co-stimulatory molecule, was decreased in CD8 T cells from LUAD. The increased expression of CD96 and decreased expression of CD226 contributed to a hypo-immune response, which impaired the antitumor immune response in LUAD. The activity of AKT-mTOR signaling was decreased in CD96+ CD8 T cells, implying the low activity of CD8 T cell function. We confirmed this by measuring cytokine production in CD8 T cells. CD96^+^ CD8 T cells expressed substantially lower levels of IFNγ and TNFα than CD96^−^ CD8 T cells did. LUAD cells might suppress the immune response by inducing the imbalance of CD96/CD226 expression on CD8 T cells in the tumor microenvironment.

The tumor microenvironment has played a critical role in shaping the immune response (Whiteside, 2008). We showed that LUAD cells induced the expression of CD96 and suppression of CD226 on CD8 T cells when they were co-cultured together. Blocking CD96 restored CD8 T cell function as inhibited by LUAD cells. LUAD cells impaired CD8 T cell function, and CD155 overexpression in cancer cells further decreased impaired CD8 T cell function. Blocking CD96 neutralized CD155-mediated inhibition of CD8 T cells. LUAD cells induced the imbalance between CD96 and CD226 expression on CD8 T cells. These data demonstrated a mechanism in which LUAD cells suppress immune response in the tumor microenvironment through CD155/CD96.

In conclusion, our findings provide insights into the mechanism that CD155 facilitates tumor growth by impairing the antitumor immune response in the tumor microenvironment through CD96. Targeting CD155/CD96 to unleash CD8 T cells in the tumor microenvironment could be a novel therapeutic alternative for LUAD patients.

## Materials and methods

### Patients

Peripheral blood and primary tumor tissue samples were collected from clinically and pathologically verified lung cancer patients at the First Affiliated Hospital, Sun Yat-sen University. Age- and sex-matched blood was collected from healthy donors. Fresh tumor tissues were collected from patients with advanced LUAD. The study was approved by the Institutional Review Board of First Affiliated Hospital, Sun Yat-sen University. All animal procedures were approved by the ethics committee of the First Affiliated Hospital, Sun Yat-sen University and performed in accordance with the guidelines provided by the National Institute of Health Guide for Care and Use of Animals. Consent forms were obtained from each patient. Demographic characteristics of the included patients were described in Supplementary Table 1.

### Cell lines and cell culture

The normal human bronchial epithelial cell line BEAS-2B was obtained from ATCC (Maryland, USA). Cell lines were authenticated by cell viability analysis, short tandem repeat (STR) profiling, and isoenzyme analysis. Cell lines were screened for mycoplasma contamination as described previously (Li *et al*, 2014). Cells were grown in RPMI 1640 medium supplemented with 10% fetal bovine serum and kept in a humidified atmosphere at 37°C with 5% CO_2._

### Western blotting

Cells were collected, and the proteins were extracted, separated by SDS-polyacrylamide gels and then electro-transferred onto polyvinylidene difluoride membranes. The membranes were washed with TBST, blocked with 10% nonfat milk in TBST and incubated with anti-CD155 (Abcam, Hong Kong) or anti-β-actin (Cell Signaling Technology, USA) primary antibodies at 4°C overnight. The membranes were then washed and incubated with horseradish peroxidase conjugated anti-rabbit IgG (Abcam, Hong Kong) at room temperature for 60 min. Signals were detected by enhanced chemiluminescence (ECL).

### Immunohistochemistry

Paraffin-embedded tissues were cut into 4-μm sections. The slides were deparaffinized, and antigen retrieval was performed. The slides were incubated with anti-CD8 (1:500) and anti-CD155 (1:100) (Abcam, Hong Kong) primary antibodies at 4°C overnight. The sections were then incubated with an HRP-conjugated secondary antibody for 1 h at room temperature. Peroxidase was visualized with 3,3’ diaminobenzidine, and the slides were counterstained with hematoxylin. For immunofluorescence staining, frozen sections were fixed with acetone for 15 min. The slides were incubated with anti-CD45 (1:200), anti-CK (1:200), and anti-CD155 (1:100) primary antibodies (Abcam, Hong Kong) at 4°C overnight and then visualized with Alexa Fluor® 488 anti-rabbit (1:200), Alexa Fluor® 546 anti-mouse (1:200) (Thermo Fisher, USA), anti-rat-TRITC (1:200) (Abcam, Cambridge, UK) secondary antibodies. Images were acquired using a fluorescence microscope (Toshiba, Japan). The number of infiltrated CD8 T cells was counted from 5 different high-power areas. Protein expression levels were evaluated semiquantitatively based on staining intensity and distribution using the immunoreactive score (IRS) as described previously (Nagata *et al*, 2004) as follows: IRS =SI (staining intensity) × PP (percentage of positive cells). The SI was determined as follows: 0, negative; 1, weak; 2, moderate; and 3, strong. The PP was defined as follows: 0, <1%; 1, 1%–10%; 2, 11%–50%; 3, 51%–80%; and 4, >80% positive cells. Ten visual fields from different areas of each tumor were used for the IRS evaluation. Negative control slides were included for each staining. An IRS score that reached 3.0 was recognized as high expression; other scores were considered low expression in this study.

### Reverse transcription-polymerase chain reaction (RT-PCR)

RNA was extracted according to the manufacturer’s instructions (Qiagen, USA). Taq DNA polymerase (Fermentas, USA) was used for cDNA synthesis. Real-time PCR was performed using SYBR Green I (Roche, USA). Amplification was performed as follows: preheating at 95°C for 10 min; denaturing at 95°C for 15 s; and annealing and extension at 65°C for 45 s for a total of 35 cycles. β-actin was used as control. The primers used are listed in Supplementary Table 2.

### Flow cytometry

Cells isolated from tumor tissues, PBMCs isolated from the matched patients with LUAD or healthy controls were stained with FITC-anti-CD4, PE-anti-CD8, APC-conjugated anti-CD96, PE-CY7-anti-Granzme B, APC-CY7-anti-Perforin, APC-anti-CD226 antibodies (Biolegend, USA). For intracellular cytokine staining, T cells (1×10^6^ cells/ml) were stimulated with 500 ng/ml PMA and 1 μg/ml ionomycin at 37°C with 5% CO2, 1 μg/ml Brefeldin for 4 h. Cells were collected and stained with BV650-conjugated anti-IFNγ antibody (Biolegend, USA). The data were acquired using a cytometer machine (BD Fortessa, USA).

### Cell isolation

Fresh tumor tissue samples were obtained from patients with LUAD who underwent surgical resection of tumor or from tumor-bearing mice. Samples were minced and digested with type I collagenase (2 mg/ml) and DNase (40 U/ml) in RPMI 1640. Cells were filtered through a cell strainer and washed with phosphate-buffered saline (PBS) twice. Peripheral blood mononuclear cells (PBMCs) from the matched patients with LUAD were isolated by density gradient centrifugation. Cells were first enriched for CD8 T cells using EasySep™ human total or naïve CD8 T Cell enrichment kits (STEMCELL Technologies, Vancouver, Canada). CD56^−^ and CD56^+^ PBMC fractions were sorted by flow cytometry. Cells were checked for purity (>97%) by flow cytometry. CD96^+^ and CD96^−^ CD8 T cells were sorted using BD influx, and the purities were checked (>95%).

### Plasmids and retroviral infection

CD155 constructs were generated by sub-cloning PCR-amplified full-length human CD155 cDNA into pcDNA3.1. Stable cell line (5×10^6^ cells) expressing CD155 was selected via treatment with 0.5 μg/ml puromycin for 10 days beginning 48 h after infection. Following selection, cancer cell lysates prepared from the pooled cell populations in sampling buffer were fractionated by sodium dodecyl sulfate-polyacrylamide gel electrophoresis (SDS–PAGE) to detect protein levels via western blotting. To deplete CD155, shRNA sequences were cloned into pGV248 to generate pGV248/CD155-shRNA (containing a green fluorescent protein reporter gene) targeting CD155. A negative vector (pGV248/control-shRNA) was similarly constructed with an unrelated shRNA sequence. DNA sequencing was used to verify all inserted sequences. Transduction efficiencies were confirmed by western blot. Transduction efficiency is measured by flow cytometry and indicated as percentage of successfully transduced GFP-positive cells.

### Co-culture

To study CD8 T cell functions affected by cancer cells, CD8 T cells (1 ×10^6^ cells/ml) were co-cultured with cancer cells (5×10^6^ cells) in which CD155 was either knocked down or overexpressed. Cells were stimulated with anti-CD3/CD28 beads and co-cultured with cancer cells in 48-well plates at a ratio of 5:1. Anti-CD3/CD28 beads were still present during T-cell/LUAD co-culture. Human CD96 blocking antibody (5 μg/ml) or isotype control (Biolegend, USA) was included in some experiments.

### Xenograft mouse model

(NOG).Cg-*Prkdc^scid^*Il2rg*^tm1Sug^*/JicCrl (NOG) mice (Weitonglihua Experimental Animal Co., Ltd, Beijing, China) are severe combined immunodeficient mice. Immune reaction of human immune cells against human tumors and the underlying mechanisms can be tested using this humanized mouse model. To better understand the immune response in the tumor microenvironment in LUAD patients, we used a patient-derived xenograft (PDX) mouse model. Freshly collected human tumor tissues (from 6 independent CD155^low^ patients) were cut into 0.5-cm^3^ pieces and subcutaneously engrafted into the NOG mice. One week later, CD56^−^ PBMC (1×10^7^ cells/ml)) from the same donors were sorted by flow cytometry and injected into the mice intraperitoneally. To study whether CD155 regulates tumor growth in vivo, PDX mice were treated with recombinant human CD155 (rCD155, 5 mg/kg, R&D, USA) 3 times a week.

To further investigate the effect of CD155 on CD8 T cell immune response in the tumor microenvironment, NOG mice received PBMC (1×10^7^ cells/ml)) from the same patients intraperitoneally for immune reconstitution. NOG mice were then subcutaneously inoculated with 5×10^6^ LUADC-Vector or LUADC-CD155 cells (from 6 independent CD155^low^ patients).

The mice were monitored every 2 days for signs of morbidity and mortality. Tumor size was measured by a caliper, and tumor volume was calculated using the formula volume as follow: (length×width^2^)×π/6. In vivo bioluminescence imaging was performed using the IVIS100 system. The Living Image acquisition and analysis software (Caliper Life Sciences) were used together as described previously (Olsen *et al*, 2017).

### Statistics

The data were expressed as the mean±SEM. Statistical analyses were performed using SPSS 16.0 (Chicago, IL, USA). The differences between groups were assessed by an unpaired, two-tailed Student’s *t* test. To adjust for multiple testing, in addition to individual p-values, we used Hochberg’s step-down method to control for a family-wise-error rate at the 0.05 level. Survival curves were plotted using the Kaplan-Meier method and compared with the log-rank test. Bivariate correlation analysis was demonstrated as Spearman’s rank correlation coefficient. Where appropriate, one-way ANOVA was used and pair-wise comparison using Tukey’s method to adjust for multiple testing was applied. Two-tailed p<0.05 was considered significant.

## The paper explained

### Problem

Recent progress in immunotherapy by blocking immune checkpoints has received great benefits and dramatically improved patient survival in the oncology field. Given about 20-30% response rates of the current immunotherapy for lung cancer, other immune-related molecules or mechanisms may be involved. Finding new checkpoint inhibitors to enhance antitumor immune response in the tumor microenvironment and further improving prognosis is attracting more and more attention in the basic research and clinic application.

### Results

In this study, we found that CD155 expression was significantly increased in tumor tissue and associated with decreased immune response, leading to poor survival in lung adenocarcinoma patients. Lung adenocarcinoma cells suppressed CD8 T cell function through CD155. Recombinant human CD155 protein inhibited immune response in the tumor microenvironment and promoted tumor growth in a patient-derived xenograft mouse model. In addition, we detected increased CD96 (co-inhibitory receptor) and decreased CD226 (co-stimulatory receptor) on CD8 T cells from lung adenocarcinoma patients. In a T cell-cancer cell co-culture system, IFNγ production in CD8 T cells was suppressed and blocking CD155/CD96 could restore IFNγ production in CD8 T cells.

### Impact

LUAD cells suppress antitumor immune response through CD155/CD96-interaction. Moreover, the inhibition effect can be reversed by CD96 blocking antibody, suggesting that CD155/CD96 can serve as a potential treatment target for LUAD

## Author contributions

**Conception and design:** Zunfu Ke, Weiling He, Hui Zhang, Shuhua Li and Yongmei Cui,

**Funding support:** Zunfu Ke

**Collection and assembly of data:** Ying Zhu, Zheng Zhu, Junfeng Zhu, Yiyan Lei, Run Lin, Di Xu, Wenting Jiang and Han Wang,

**Data analysis and interpretation:** Zunfu Ke, Weiling He and Hui Zhang

**Manuscript writing:** Zunfu Ke, Weiling He and Ying Zhu

**Final approval of manuscript:** All authors.

